# *De novo* assembly and annotation of the Patagonian toothfish (*Dissostichus eleginoides*) genome

**DOI:** 10.1101/2022.12.15.520537

**Authors:** David Ryder, David Stone, Diana Minardi, Ainsley Riley, Justin Avant, Lisa Cross, Marta Soeffker, Deborah Davidson, Andrew Newman, Peter Thomson, Chris Darby, Ronny van Aerle

**Author notes:** Corresponding Author: David Ryder. Joint senior authors (Chris Darby, Ronny van Aerle).

## Abstract

Patagonian toothfish (*Dissostichus eleginoides*) is an economically and ecologically important fish species in the family Nototheniidae, found at depths between 70 and 2,500 meters on the southern shelves and slopes around the sub-Antarctic islands of the Southern Ocean. Genomic sequence data for this species is limited. Here, we report a high-quality assembly and annotation of the *D. eleginoides* genome, generated using a combination of Illumina, PacBio and Omni-C sequencing technologies. To aid the genome annotation, the transcriptome derived from a variety of toothfish tissues was also generated using both short and long read sequencing methods. The final genome assembly was 797.8 Mb with a N50 scaffold length of 3.5 Mb. Approximately 31.7% of the genome consisted of repetitive elements. A total of 35,543 putative protein-coding regions were identified, of which 50% have been functionally annotated. Transcriptomics analysis showed that approximately 64% of the predicted genes (22,617 genes) were found to be expressed in the tissues sampled. Comparative genomics analysis revealed that the anti-freeze glycoprotein (AFGP) locus of *D. eleginoides* does not contain any AFGP proteins compared to the same locus in the Antarctic toothfish (*Dissostichus mawsoni*). This is in agreement with previously published results looking at hybridization signals and confirms that Patagonian toothfish do not possess AFGP coding sequences in their genome. The high-quality genome assembly of the Patagonian toothfish will provide a valuable genetic resource for ecological and evolutionary studies on this and other closely related species.

## Introduction

Patagonian toothfish (*Dissostichus eleginoides*) are found at depths between 70 and 2,500 m, on the southern shelves and slopes around the sub-Antarctic islands of the Southern Ocean (Collins *et al*., 2010; Canales-Aguirre *et al*., 2018a). *D. eleginoides* and its closely related sister species, the Antarctic toothfish (*Dissostichus mawsoni*), are both within the Nototheniidae clade, and their geographical distributions, for the most part, do not appear to overlap (Soeffker *et al*., 2022, Roberts, Xavier and Agnew, 2011).

The two toothfish lineages have separated, with one fish species, the Patagonian toothfish, adapting itself to more temperate climates, and Antarctic toothfish retaining all the genetic adaptations required to survive in the cold Antarctic waters (Colombo *et al*., 2015).

There are some large differences between the two species which is illustrated by the fact that, in contrast to the Antarctic toothfish, no evidence has been found for the presence of anti-freeze glycoprotein (AFGP) in the blood of the Patagonian toothfish (Gon and Heemstra, 1990) or within its genome (Christina Cheng and William Detrich, 2007). Whilst the region representing the AFGP locus in the Antarctic toothfish genome was fully sequenced using Sanger sequencing (BAC clones) by Nicodemus-Johnson *et al*. (2011) this paper is the first to try and characterise the AFGP locus in Patagonian toothfish.

Other, more subtle adaptations of Nototheniidae fish species to the Antarctic environment include changes in membrane composition and the structure of protein translocation channels (Römisch *et al*., 2003; Bargelloni *et al*., 2019), in the regulation of molecular chaperones (Hofmann *et al*., 2000; Place and Hofmann, 2005; Bilyk and DeVries, 2011; Bilyk and Cheng, 2013, 2014), in the expression of haemoglobin and regulation of the circadian rhythm (B. M. Kim *et al*., 2019), as well as in the structure of microtubules in the cytoplasm (Detrich *et al*., 2000), though more work is required to obtain a complete picture of all the different ways in which this group has adapted to such cold conditions.

Recent advances in sequencing technologies have facilitated the sequencing of genomes of any species of interest in a relatively cost-effective way. Availability of genomic sequence data facilitates studies on many aspects of fish biology, including the evolution of adaptation to cold water conditions. In 2019, the Antarctic toothfish genome was sequenced (Chen *et al*., 2019) and more recently, the genomes of 24 different notothenioid fish species were sequenced and assembled (Bista *et al*., 2022). Despite supporting valuable fisheries and the important ecological role it plays in the Southern Ocean ecosystem, the genome of Patagonian toothfish had not been sequenced and very limited genomic sequence information available in the public databases for this species. This lack of sequence information restricts studies on for example physiology and disease resistance in this species, but also on population genetics and comparative genomics analysis with other (related) fish species.

The aim of this study was to generate a high-quality genome assembly of the Patagonian toothfish using a combination of long read (PacBio), short read (Illumina) and Omni-C sequencing data. The genome sequence was characterised and screened for the presence of the AFGP locus and/or AFGP genes and subjected to phylogenetic analysis with related fish species. The annotated Patagonian toothfish genome generated in this project provides a valuable genetic resource for studying the evolution of adaptation of fish to cold waters, as well as other evolutionary and ecological studies on this species.

## Methods

### Sample collection

A mature female Patagonian toothfish (*D. eleginoides*) (95 cm in length, 8.18 kg in weight) was caught between 1300 and 1350 m in depth in a fishing area near South Georgia (CCAMLR subarea 48.3B). Samples from the spleen, muscle, liver, kidney, intestine, heart, ovary, gills and brain were collected and preserved in RNAlater™ (Invitrogen).

### DNA extraction

High molecular weight DNA was extracted from pooled visceral tissues (spleen, liver, and kidney) stored in RNAlater™ using the Qiagen Genomic tip 500G and the manufacturer’s recommended protocol. Briefly, 400 mg of toothfish visceral tissues were ground to a powder with liquid nitrogen in a pre-cooled mortar and pestle and then digested in 20 ml of G2 buffer containing RNase and proteinase K for 2 h at 50 °C. Following digestion, the sample was loaded onto a pre-equilibrated genomic tip 500G. The column was washed twice with 15 ml Buffer QC and eluted 15 ml buffer QF. The DNA was precipitated using 0.7 volumes (10.5 ml) of isopropanol and centrifugation at 5000 g for 15min at 4 °C; the pellet was washed with 4 ml of cold 70% ethanol and centrifuged at 5000 g for a further 10 min at 4 °C. The DNA pellet was air dried for 10 min and resuspended in 500 µl of TE buffer and stored at –80 °C.

### RNA extraction

RNA was extracted from the nine individual tissue samples stored in RNAlater™ using the Direct-zol™ RNA Miniprep Plus kit (Zymo Research) according to the manufacturer’s instructions. Briefly, 50 mg of each tissue were ground to a powder with liquid nitrogen in a precooled mortar and pestle and lysed in 600 µl of TRI Reagent. The lysed samples were transferred to clean 1.5 microtubes and equal volumes of absolute ethanol added. The samples were thoroughly mixed and 700 µl transferred to Zymo-Spin IIICG columns and centrifuged at 13,000 g for 30 sec. The flowthroughs were discarded, and the columns treated with DNase I prior to washing with 400 µl of Direct-zol RNA prewash, followed by a wash with 700 µl of RNA wash buffer. The RNA samples were then eluted in 100 µl DNase/RNase free water and stored at -80 °C.

### Genome Sequencing

Sequence library preparation (except for the Omni-C library) and subsequent sequencing were conducted by the Exeter Sequencing Facility (University of Exeter). Genomic DNA libraries were generated using the SMRTbell Express Template Prep Kit 2.0, following the protocol described online (PacBio, 2019). An additional enzymatic digest step (to remove any linear molecules) was included as described on page 13 of the protocol before size-selection of >20 kb fragments on a high pass cassette on the Blue Pippin (Sage Science, MA, USA). A total of 6 SMRT cells were run on the PacBio Sequel.

An Illumina sequencing DNA library was prepared from the same DNA sample using the NEBNext® Ultra™ II FS DNA Library Prep Kit for Illumina (New England Biolabs). An Omni-C library was prepared for a separate visceral sample using the Dovetail™ Omni-C™ Kit with Library Module and Primer Set for Illumina (Dovetail Genomics, CA, USA) according to the manufacturer’s protocol. Both sequencing libraries were run using an Illumina SP flow cell on an Illumina NovaSeq (2×150 bp protocol), following manufacturer’s instructions (Illumina, San Diego, CA, USA).

### RNA Sequencing

RNA quality was assessed with an Agilent TapeStation using the RNA Analysis ScreenTape System (Agilent). Extracted RNA from brain, muscle, gills, and kidney passed the RIN score threshold required for IsoSeq sequencing. Equimolar amounts of RNA from these samples were pooled and used to generate a single SMRTBell library, as described in a paper by Leung *et al*. (2021) and using version 3 chemistry, and equal amounts of the library were run over 3 SMRT cells on the PacBio Sequel. For short read sequencing, mRNA libraries were prepared for all nine tissue samples using the TruSeq Stranded mRNA sample preparation kit (Illumina) and run on an Illumina NovaSeq (2x 150 bp protocol).

### Assembly of Genome

The size of the genome was estimated by using Jellyfish 2.2.10 to count the frequency of each 21-mer in the Illumina reads (Marçais and Kingsford, 2011), using the same software to create a histogram for every k-mer whose frequency was between 1 and a million, and subsequently importing the results into GenomeScope Release 1 (Vurture *et al*., 2017).

Canu version 2.2 was used to carry out an initial assembly using the default assembly parameters provided for PacBio reads (Koren *et al*., 2017), with the estimated genome size being set to 800 Mb.

The Illumina reads were mapped to the assembly using minimap 2.20-r1061 and the default alignment parameters for short paired-end reads (Li, 2018, 2021). Alignment results were sorted and indexed using Samtools 1.12 (Li *et al*., 2009). Pilon 1.24 was used to polish the assembly, specifically being configured to identify and fix any single nucleotide polymorphisms (SNPs) and indels identified in the alignment, as well as the ‘diploid’ option being set to try and improve calling of heterozygous SNPs.

The PacBio reads were mapped to the assembly using minimap 2.20-r1061 and the default alignment parameters for PacBio Continuous Long Reads (CLR) reads. The maximum size of the index used during alignment was set to 1600G, with secondary alignments being excluded from the final output. Alignment results were sorted and indexed using Samtools 1.12 (Li *et al*., 2009). Purge_haplotigs 1.1.2 was used to plot a histogram showing coverage of the genome (Roach, Schmidt and Borneman, 2018). Based on coverage of the genome the low, middle and high coverage thresholds were manually set to 10, 65 and 150, respectively, which were then subsequently used to calculate coverage statistics for each contig and flag suspicious contigs both using the ‘cov’ command in purge_haplotigs. Finally, the assembly was separated out into primary contigs, haplotigs and artefacts using the purging algorithm. Unless otherwise specified, all the steps run using purge_haplotigs were done using default parameters.

The Omni-C sequences were mapped to the assembled primary contigs using bwa 0.7.17-r1188 (Li, 2013) and subsequently filtered based on the Omni-C protocol published online by Dovetail Genomics (Dovetail Genomics, 2021). Briefly, pairtools 0.3.0 was used to find ligation junctions and remove PCR duplicates (Open Chromosome Collective, 2019), with Samtools 1.15 being subsequently used to sort the results. The parameters and exact commands used during this stage in the analysis were as described in the protocol online. The bamToBed script included with bedtools 2.30.0 was then used to convert any alignments identified within ligation junctions from the bam to bed file format (Quinlan and Hall, 2010), with the bed file subsequently being sorted based on read name using the sort command (which is included as part of GNU’s coreutils package). SALSA 2.3 was used to scaffold the assembled primary contigs using the Omni-C alignments processed and filtered as described above (Ghurye *et al*., 2019), with default parameters being used, the enzyme being set to ‘DNASE’ and the option for finding misassemblies being enabled. Following scaffolding of the primary contigs, alignment of the PacBio reads to the scaffolds and purging of haplotigs was repeated, this time using a lower ‘middle coverage’ threshold of 62, with all the other parameters kept the same.

Meryl 1.3 was used to count 21-mer k-mers in the Illumina reads (Maryland Bioinformatics Labs, 2021). Merqury 1.3 was used to compare k-mer counts from the Illumina reads against different polished and unpolished versions of the assembly (Rhie *et al*., 2020), to allow a more accurate comparison of how different steps in the analysis impacted the quality and completeness of the assembly. BUSCO 4.1.2 was used to assess the level of completeness of the genome by comparison of predicted proteins or transcripts against v10 of the OrthoDB for species in the Actinopterygii lineage (Kriventseva *et al*., 2019; Manni, Matthew R. Berkeley, *et al*., 2021; Manni, Matthew R Berkeley, *et al*., 2021).

### Processing and mapping of IsoSeq3 RNA Sequencing data

Subreads were used to output circular consensus sequences with a minimum predicted accuracy of 0.9 using the ccs v4.2.0 software. Barcodes were trimmed from circular consensus sequences using lima v1.11.0, which was parameterised using the IsoSeq preset, the peek guess option, and with the sequence of primers being based on those that would be expected when using a Clontech SMARTer and NEB cDNA library preparation method. The trimming of poly-adenosine tails and removal of concatemers was carried out using the refine command in isoseq3 v3.3.0. The full length non-chimeric reads were then converted from the bam to fastq format using bam2fastq v1.3.0. The software used for initial processing of IsoSeq3 data was from the official release of SMRT Link version 9.0.0 (Pacific Biosciences of California, 2020).

The full length non-chimeric long reads sequenced and initially processed using the IsoSeq3 protocol were aligned against the draft genome using the splice:hq preset in minimap 2.20 (Li, 2018, 2021), which was configured to find conical splicing sites on only the forward transcript strand, with samtools 1.15 being subsequently used to sort the results. StringTie 2.1.4 was then used to assemble reads into potential transcripts using the long read processing option (Kovaka *et al*., 2019).

### Processing and mapping of Illumina RNA Sequencing data

Short read, paired end sequencing data from each sample were separately aligned against the draft genome using hisat 2.2.1 with default options, except for the rna-strandness option, which was set to ‘RF’, and with reported alignments tailored specifically for transcript assemblers (D. Kim *et al*., 2019). Samtools 1.15 was used to sort the results. StringTie 2.1.4 was used to assemble reads of each individual sample into potential transcripts using default parameters (Kovaka *et al*., 2019).

### Annotation of Genome

RepeatModeler 2.0.1 was used for *de novo* detection of repeats and the creation of a custom repeat library with the long terminal repeat (LTR) structural discovery pipeline enabled (Flynn *et al*., 2020), with the maximum sample size to be used by RECON set to 81 Mb, with relevant dependencies including TRF 4.09 (Benson, 1999), RECON 1.08 (Bao and Eddy, 2002), RepeatScout 1.0.6 (Price, Jones and Pevzner, 2005) and LTR_Retriever 2.9.0 (Ou and Jiang, 2018). RepeatMasker 4.1.1 was then used to identify and soft mask repeats in the assembly (Smit, Hudley and Green, 2013), with RMBlast 2.10.0 acting as a search engine (Smit, Hudley and Rosen, 2019) and a minimum alignment score of 250 being used to mask repeats.

StringTie 2.1.4 was used to merge transcripts identified from the different sequencing libraries, as described above, with the stringtie2gff3 utility from Funannotate 1.8.9 then being used to convert the predicted gene structures from GTF to GFF3 format (Kovaka *et al*., 2019). Funannotate 1.8.9 was used to predict gene coding regions with the type of organism set to ‘other’, the maximum intron length set to 500,000 bp, repeat regions being used in EVM consensus model building, OrthoDB v10 for species in the Actinopterygii lineage being used as the BUSCO database, the predicted gene structures from StringTie2 being provided as a set of pre-computed transcript alignments to be used in training gene model prediction algorithms and with proteins from the *Trematomus bernacchii* (NCBI Genbank Accession GCF_902827165.1), *Cottoperca gobio* (GCF_900634415.1), *Notothenia coriiceps* (GCF_000735185.1), and *Pseudochaenichthys georgianus* (GCF_902827115.1) genome assemblies mapped to the genome and used to provide evidence with which gene coding regions could be identified.

InterProScan 5.56-89.0 was used to carry out functional annotation of the genome using all default analysis modules (Jones *et al*., 2014; Blum *et al*., 2021). Funannotate 1.8.9 was used to provide functional annotation of gene coding regions within the genome, incorporating InterPro and Gene Ontology (GO) terms based on InterProScan results, gene and product names based on a BlastP search of predicted proteins against UniProt DB 2022_01 (Camacho *et al*., 2009; The UniProt Consortium, 2019), and additional annotations from Pfam-A, MEROPS (Rawlings, Barrett and Bateman, 2010), CAZYme (Drula *et al*., 2022), BUSCO2 (Manni, Matthew R. Berkeley, *et al*., 2021; Manni, Matthew R Berkeley, *et al*., 2021), and Phobius analyses (Käll, Krogh and Sonnhammer, 2004).

### Phylogenetic Analysis

A range of species (n = 32) from within and outside of the Notothenioid taxonomic clade (n = 10) were identified for which whole genome sequencing data was available in public NCBI databases (see Supplemental Table 1 for full list of species). BUSCO 4.1.2 was run in genome mode against assembled genomes from each of the different species (Manni, Matthew R. Berkeley, *et al*., 2021; Manni, Matthew R Berkeley, *et al*., 2021), with Augustus 3.3.3 being used for prediction of gene coding regions (Stanke *et al*., 2006), and OrthoDB v10 for species in the Actinopterygii lineage as a reference (Kriventseva *et al*., 2019). A custom python script was then run against the output of BUSCO to identify single copy orthologs which were consistently observed across every species, with the results being grouped by ortholog in such a way that a set of gene coding regions was available for each ortholog, which summarised all the relevant sequences observed across the different species. The trimNonHomologusFragments function from MACSE 2.05 was then run with default parameters to remove non-homologous sequences from the 5’ and 3’ ends of each sequence from each species assigned to a given ortholog (Ranwez *et al*., 2011). Sequences corresponding to each ortholog were aligned using the alignSequences function from MACSE 2.05. Translated amino acid sequences were checked by HmmCleaner 0.180750, with any regions thought to be due to sequencing error, rather than biological variation, being masked. The reportMaskAA2NT function from MACSE 2.05 was used to mask any regions identified as problematic by HmmCleaner in the corresponding nucleotide alignment, as well as some additional processing on the aligned sequences (parameters used: -min_NT_to_keep_seq 30 -min_seq_to_keep_site 4 -min_percent_NT_at_ends 0.9 -dist_isolate_AA 3 - min_homology_to_keep_seq 0.3). Following this process trimmed, aligned, homologous sequences longer than 500 bp were available for 151 orthologs from 42 different species, making up a total of 220,443 bp of sequencing data.

IQ-TREE 2.2.0.3 was used to carry out a partitioned, maximum likelihood analysis on each of the 151 nucleotide alignments (Minh *et al*., 2020), with each partition sharing the same set of branch lengths, but allowing different rates of evolution, using the relaxed hierarchical clustering algorithm to examine the top 10% of partition merging schemes and identifying the best option (Chernomor, von Haeseler and Minh, 2016), using an ultrafast bootstrapping approach with 1000 replicates (Hoang *et al*., 2018), and identifying the best-fit substitution model following identification of the best partitioning scheme (Kalyaanamoorthy *et al*., 2017).

### Comparative Genetics

A range of species from within (n = 9) and outside of the *Notothenioid* taxonomic clade (n = 5) were identified for which whole genome sequencing data and accompanying gene annotations were available. OrthoFinder 2.5.4 was used to identify orthologs present across the different species (Emms and Kelly, 2015), as well as provide a phylogenetic tree based on common orthologs shared across the different taxa, calculate various statistics, and identify gene duplication events.

### Assembly of Mitochondrial genome

The mitochondrial genome was assembled from the Illumina reads using GetOrganelle 1.7.5 (Jin *et al*., 2020) and the following parameters: -F animal_mt -R 15 --target-genome-size 19000.

Since the initial mitochondrial genome included a few unexpected repeats and pseudogenes the PacBio reads was mapped to the initial assembly using minimap 2.20-r1061 and the default alignment parameters for PacBio CLR reads. Samtools 1.15 was used to filter the results to exclude any unmapped alignments, with a list of reads which mapped against the initial draft assembly then being output to a text file. Seqtk 1.3-r106 was used to create fastq files for each of the PacBio SMRT cells which included reads that mapped against the mitochondrial genome. Canu 2.2 was then used to assemble the mitochondrial genome with default assembly parameters provided for PacBio reads (Koren *et al*., 2017), the estimated genome size being set to 20 kb and with the ‘corOutCoverage’ parameter set to 10,000x coverage.

To polish the assembly based on long reads, Illumina reads were mapped to the assembly using minimap 2.20-r1061 and the default alignment parameters for short paired-end reads (Li, 2018, 2021). Alignment results were sorted and indexed using samtools 1.12 (Li *et al*., 2009). Pilon 1.24 was used to polish the assembly, specifically being configured to identify and fix any SNPs and indels identified in the alignment, as well as the ‘diploid’ option being set to improve calling of heterozygous SNPs.

Following assembly of the mitochondrial genome using long reads and subsequent polishing of the genome using Pilon and short reads, the genome was annotated with the mitos2 webserver (Donath *et al*., 2019).

### Identifying differentially expressed transcripts

Indexes to be used by the STAR alignment algorithm (Dobin *et al*., 2013) were generated from the reference transcriptome (consisting of all predicted genes by Funannotate) using the rsem-prepare-reference command from RSEM 1.3.1 (Li and Dewey, 2011). The rsem-calculate-expression command was used to align short read, paired end reads from each sample against the reference transcriptome and calculate expected gene expression levels, with the strandedness option set to reverse and STAR 2.7.10a being used for alignment (Dobin *et al*., 2013). The rsem-generate-data-matrix command was then used to convert the expected gene expression levels into a format more suitable for downstream analysis. Since each tissue sample was only sequenced once, with no treatments being available whose impact could be investigated, it was decided to compare gene expression levels in each tissue against other tissues, to detect gene expression level variations across different tissues. For each tissue, expected gene expression levels were imported into R 4.1.2 (R Core Team, 2021) and the DGEList function was used to convert the results into a format suitable for use with the edgeR 3.36.0 package (Robinson, McCarthy and Smyth, 2010). Low abundance transcripts were filtered out of the dataset by running the filterByExpr function with default parameters. The calcNormFactors function was used to calculate data scaling factors for the different libraries. Differentially expressed genes between two experimental groups (each tissue vs all other tissues pooled together) were identified using the exactTest function, with the square root dispersion value set to 0.4, since there were no biological replicates. The gene lists were then sorted (ranked) by the log2-fold change in gene expression between the two experimental groups and subsequently used to carry out a gene set enrichment analysis (GSEA) of gene ontology using the gseGo function from the clusterProfiler 4.2.2 package (Wu *et al*., 2021). For the GSEA, default parameters were used, as well as a custom database with gene ontologies inferred using InterProScan and Funannotate, as described above, and converted into an appropriate format using the AnnotationForge 1.36.0 package (Carlson and Pagès, 2021). Transcripts were considered to be expressed in a tissue when expected read counts were ≥ 10 (as determined by RSEM).

### Identifying/characterising the AFGP Locus

The sequence and gene annotations for one of the published haplotypes of the AFGP locus in Antarctic toothfish was downloaded from NCBI in fasta and gff format (NCBI Genbank Accession HQ447059.1). The sequence of each transcript predicted for the AFGP locus was then output in fasta format using gffread 0.12.7 (Pertea and Pertea, 2020). The full length, non-chimeric, long reads sequenced and initially processed using the IsoSeq3 protocol were aligned against predicted transcripts for the AFGP locus using the map-hifi preset in minimap 2.20 (Li, 2018, 2021). Short read, paired end reads from the transcriptome of each sample were also aligned against predicted transcripts for the AFGP locus using the default alignment parameters for short, paired end reads in minimap 2.20. Alignment results were then subsequently sorted and various coverage statistics were calculated using Samtools 1.15 to determine which transcripts in the AFGP locus for Antarctic toothfish were being actively expressed within Patagonian toothfish.

The draft genome assembly, including both primary contigs and haplotigs, was aligned against both of the publicly available haplotypes for the AFGP locus in Antarctic toothfish (NCBI Genbank Accessions HQ447059.1 and HQ447060.1) using nucmer 4.0.0rc1 (Marçais *et al*., 2018). Alignments between the draft genome and the AFGP locus were checked to identify any alignments which had more than 90% identity and which were longer than 1000bp, with candidate sequences filtered based on the part of the AFGP locus with which they shared a high percentage identity, as well as the overall proportion of the AFGP locus which was matched. Regions of interest were then checked for the presence of *hsl* and *tomm40* genes, which have previously been identified as being situated at the 5’ and 3’ end of the AFGP locus (Bista *et al*., 2022). These genes were also cross referenced against results from the OrthoFinder analysis, to check the genes in question were orthologs, and not paralogs, as well as confirm that corresponding sequences in the assembled genomes of species such as *C. gobio* and *P. georgianus* included similar genetic elements such as trypsin, peptidase and AFGP genes.

The PacBio reads were mapped to the candidate AFGP loci using minimap 2.20-r1061 and the default alignment parameters for PacBio CLR reads. Samtools 1.12 was used to sort and index the results and to exclude alignments which were less than 10 kb in length relative to the reference.

Blastn was used to identify regions of similarity between the AFGP loci of Patagonian toothfish (scaffold_69; position 226,982-461,816) and Antarctic toothfish (HQ447059.1; position 516-438,650). A schematic representation showing the annotated AFGP loci of the two species of toothfish alongside that of the same locus in the *C. gobio* genome (NC_041370.1; position 3,945,475-4,065,448) was created using gggenes v0.4.1 (https://wilkox.org/gggenes/).

## Results

### Assembly and Scaffolding of the Genome

Long read sequencing using 6 SMRT cells generated a total of 77.1 gigabases (Gb), with an average of 12.8 Gb per SMRT cell. The longest read generated was 130 kb, with the N90 and N50 values being 10 and 32 kb respectively. Illumina sequencing of the same DNA sample produced approximately 979 million read pairs (148 Gbp).

Based on the Illumina sequencing library and a k-mer based statistical approach, the genome size was estimated to be approximately 762 Mb, with 242 Mb of repeats. The initial genome assembly, after polishing with Illumina reads and purging haplotigs from the assembly was 799 Mb, consisting of 593 contigs and with N50 values of 2.54 Mb. Following assembly of the genome, the sequences derived from the Omni-C library were mapped to the assembled contigs. Table 1 shows a summary of the proportion of mapped and unmapped reads, the number of PCR duplicates as well as alignment results sorted into cis and trans pairs and estimated insert lengths for cis pairs. Initial results suggested a high proportion of unmapped or partially mapped read pairs in the library and relatively short distance interactions in those cases where both reads in a pair mapped to the same contig, suggesting the library may have had some contamination, there was insufficient DNA, or DNA was of insufficient quality prior to starting the Omni-C protocol. Despite this, when using the results to scaffold the genome with SALSA, the number of contigs in the genome assembly was reduced from 593 to 455, with the median contig length increasing from 2.54 to 3.55 Mb. Following scaffolding of the assembly it was purged of any remaining haplotigs using stringent coverage thresholds, resulting in a final genome assembly of 448 contigs totalling 797.8 Mb (Table 1 and Figure 1) of which 253 Mb (31.76 %) consisted of repeat elements (see Supplemental Table 2). Overall, including haplotigs, a k-mer based analysis of genome completeness using Merqury suggested the assembly was 97.62 percent complete. As part of the same analysis, the consensus quality value (QV) of the assembled, scaffolded, purged assembly was calculated as 42.09 (see Table 2).

**Table 1:** Results from evaluating the quality of the Omni-C library including (a) statistics on the initial alignment as well as subsequent filtering of low-quality alignments, PCR duplicates and unpaired reads and (b) classifying read pairs, characterising the insert size and identifying valid pairs.

|  | Number of reads | Percentage of reads |
| --- | --- | --- |
| Paired reads | 329,516,656 | 100.00% |
| Unmapped read pairs | 55,047,299 | 16.71% |
| Mapped, unpaired reads | 124,028,116 | 37.64% |
| Mapped, paired reads with PCR duplication | 85,290,350 | 25.88% |
| Mapped, paired reads with no PCR duplication | 65,150,841 | 19.77% |

(b)
|  | Number of reads | Percentage of mapped paired reads (no PCR duplication) |
| --- | --- | --- |
| Cis Read Pairs | 18,793,875 | 28.85% |
| Trans Read Pairs | 46,356,966 | 71.15% |
| Valid Read Pairs (cis ≥ 1kb + trans) | 48,308,460 | 74.15% |
| Cis Read Pairs < 1kb | 16,842,381 | 25.85% |
| Cis Read Pairs ≥ 1kb | 1,951,494 | 3.00% |
| Cis Read Pairs ≥ 10kb | 1,420,337 | 2.18% |

**Table 2:** Statistics summarising the contiguity, quality, and completeness of the Patagonian toothfish (*Dissostichus eleginoides*) genome assembly. The consensus quality and k-mer completeness statistics were calculated by comparing the genome to short reads using Merqury.

|  | Primary Contigs | Haplotigs | Overall |
| --- | --- | --- | --- |
| Total Size (Mb) | 798 | 651 | 1,449 |
| Scaffold N50 Length (kb) | 3,550 | 201 | 1,247 |
| Scaffold N90 Length (kb) | 900 | 103 | 125 |
| Number of Contigs | 448 | 3,524 | 3,972 |
| QV score | 42.09 | 37.99 | 39.78 |
| Completeness score | 92.51 | 68.31 | 97.62 |

**Figure 1:**
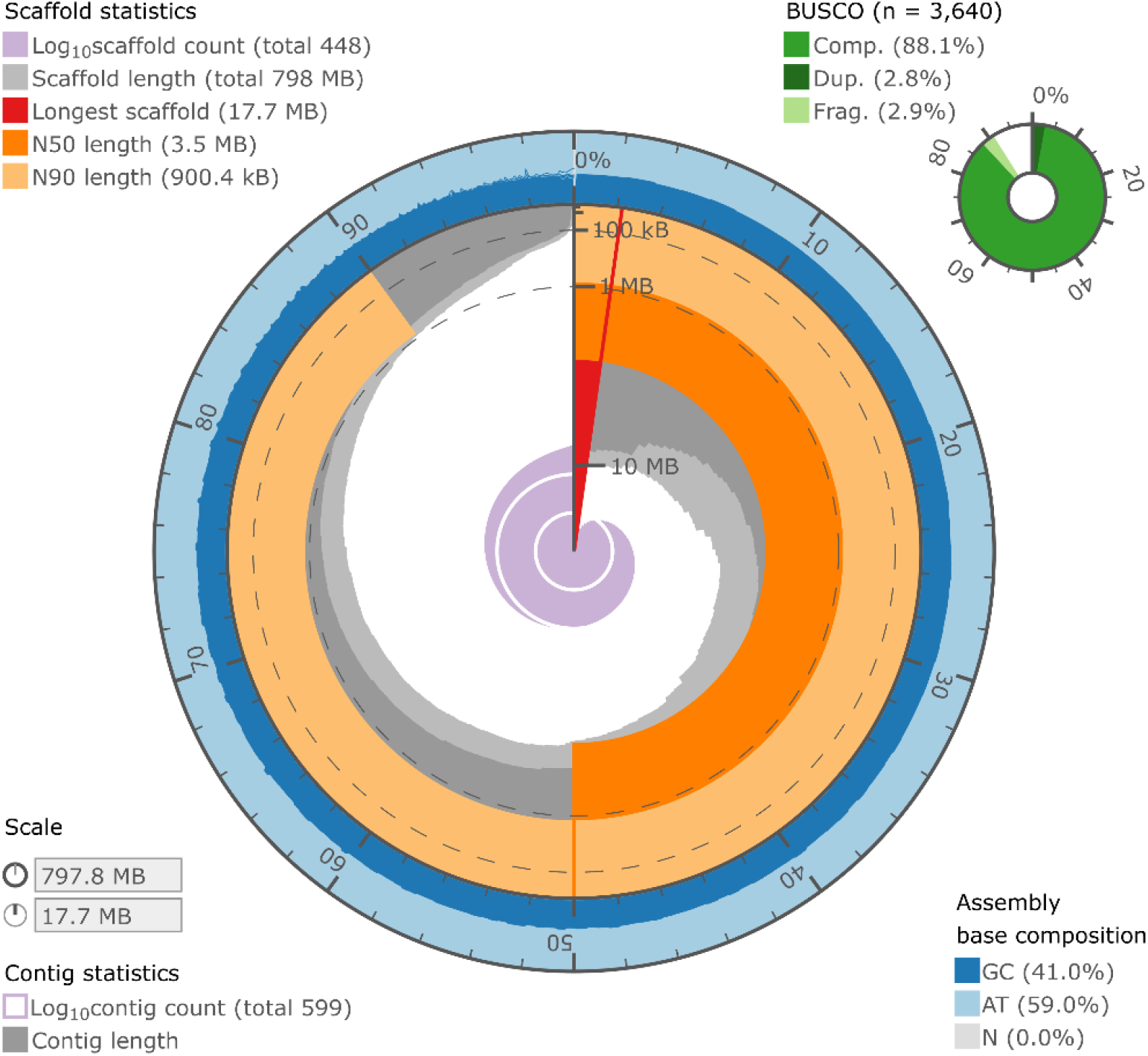
Visualisation of the Patagonian toothfish (*Dissostichus eleginoides*) genome assembly statistics using assembly-stats (https://github.com/rjchallis/assembly-stat). Red, dark and light orange represent the longest, N50 and N90 scaffold lengths, respectively, whilst the dark grey colour represents the length of each scaffold plotted against the cumulative length of all scaffolds on a circular axis, with the longest scaffold plotted nearest the red marker and scaffolds being sorted according to length. The light grey colour follows the same approach, but for contigs, rather than scaffolds. The outermost layer, plotted in blue, shows the GC content across the genome. The BUSCO score was determined by running BUSCO against the final set of gene annotations using ‘proteins’ mode and the OrthoDB v10 *Actinopterygii* database.

### Transcript assembly, analysis and genome annotation

RepeatMasker identified 253.40 Mb of repeats which were soft masked prior to genome annotation and used by EVidence Modeler (EVM) whilst annotating genes. Augustus, SNAP and GlimmerHMM were used to carry out *ab initio* gene prediction, with all three using BUSCO predicted genes as training data, but with Augustus also using results from the mapping of reference proteomes and RNA sequencing data to the genome as *‘hints’*. In total there were 35,543 predicted protein-coding sequences after combining the various predictions using EVM, as well as 6,887 predicted tRNA sequences. Predicted protein coding regions in the final set of gene annotations included complete copies of more than 90% of the proteins present within the OrthoDB v10 *Actinopterygii* database, according to BUSCO. Functional annotation of the genome included adding 51,105 gene ontology terms, 4,438 signal peptide predictions and 7,453 transmembrane annotations.

A total of 22,617 different genes were expressed in at least one of the tissues, of which 3,762 genes were expressed in every tissue (Table 3). Each tissue had uniquely expressed genes, with the brain (n = 358) and ovary (n = 123) having the most, whilst the spleen did not express any unique genes. The top 10 biological processes that were found to be enriched in each of the tissues are shown in Figure 2 (see Supplemental Table 3 for complete lists of enriched gene ontology terms and corresponding expressed genes for each tissue). As expected, the processes identified to be enriched in each of the tissues related to the specific function(s) of those tissues. For example, the brain, which had the highest number of uniquely expressed transcripts, was enriched for biological processes relating to signalling across synapses, neurotransmitter secretion and central nervous system development. Some tissues, such as the intestine, did not have any biological processes which were identified as enriched, whilst the spleen only had one, which was *‘protein localisation to the endoplasmic reticulum’*. The small number of enriched terms for the spleen could possibly be due to overlapping functions with other tissues.

**Table 1:**
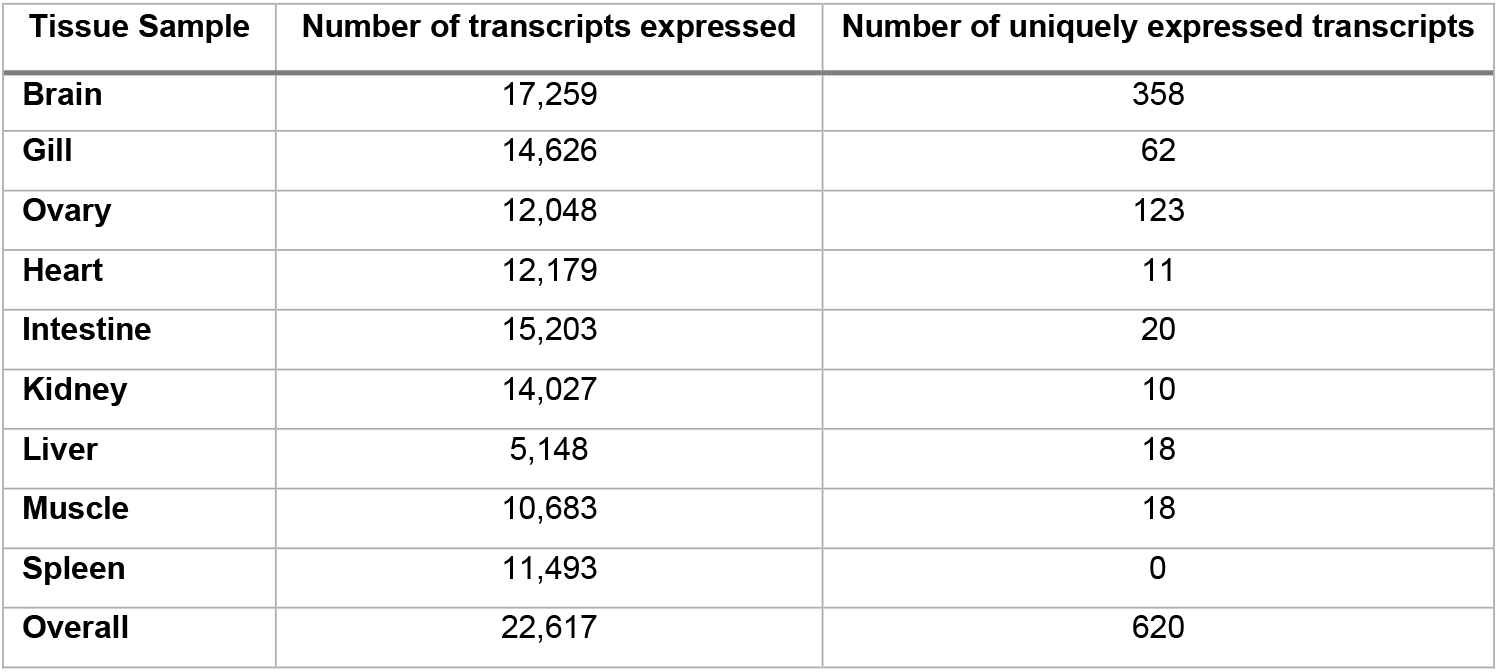
The number of transcripts (total and unique) expressed in each tissue sample.

**Figure 2:**
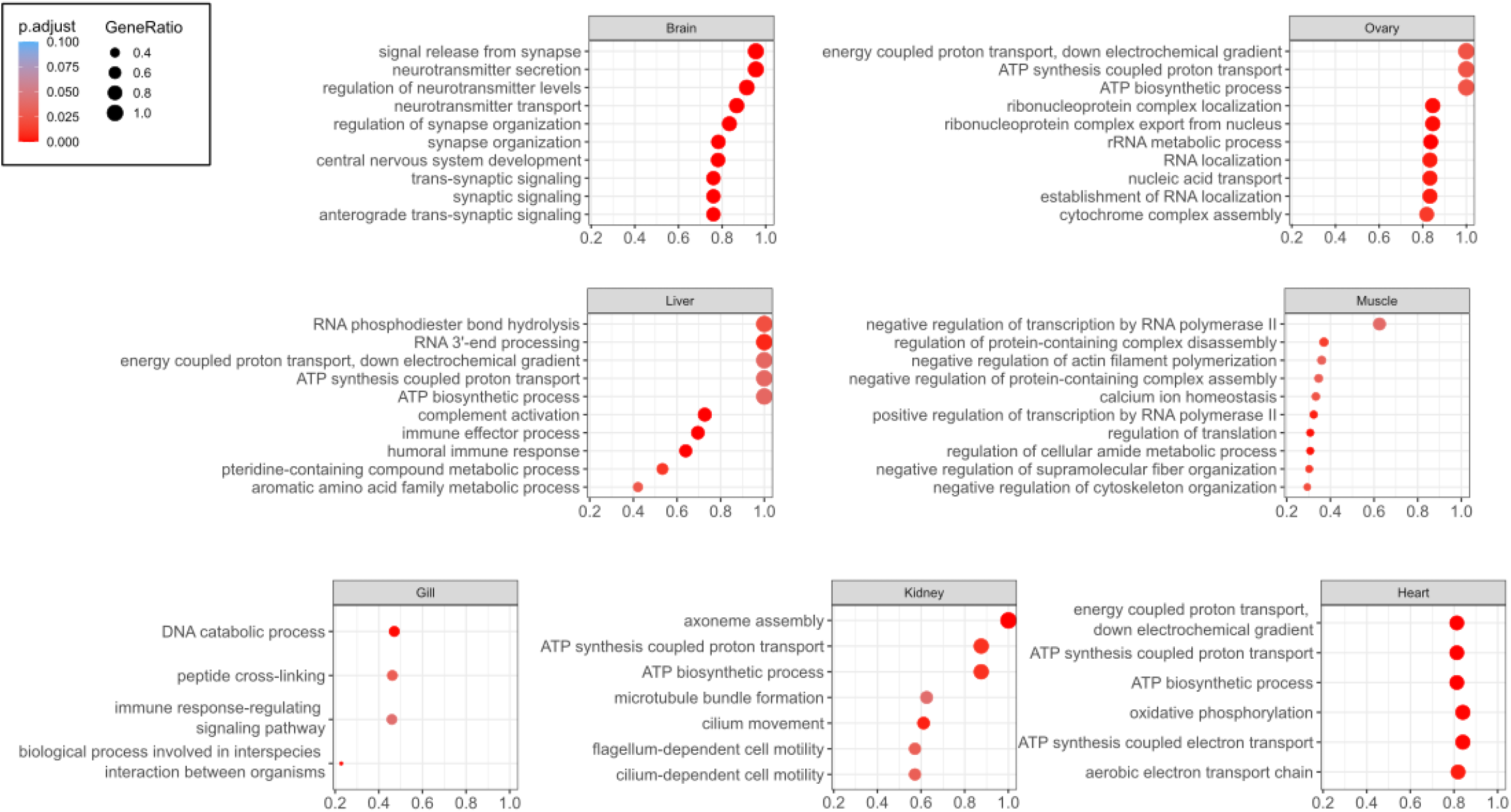
Dot plots showing the top 10 (or all) enriched gene ontologies (Biological Processes) as identified by Gene Set Enrichment Analysis (GSEA) in each tissue. The size of the solid circles is proportionate to the number of genes represented in the corresponding category and the colour indicates the significance value.

### Phylogenetic analysis

Bootstrap analyses provided high confidence for the maximum likelihood phylogenetic tree topology generated based on an alignment of 220 kb of DNA sequences from 151 orthologs and 42 percomorph fish species (Figure 3). There were however a few exceptions, including locations within some of the deeper branches of the tree, outside of the notothenioid clade of fish species, as well as for some *Trematomus* and Artedidraconidae species. In the latter case, poor support for placement of individual species within these taxonomic groups did not seem to impact overall placement of the *Trematomus* genus or the Artedidraconidae family within the phylogeny tree. The Patagonian toothfish sequences branched with the Antarctic toothfish near the root of the Antarctic notothenioid spp. clade.

**Figure 3:**
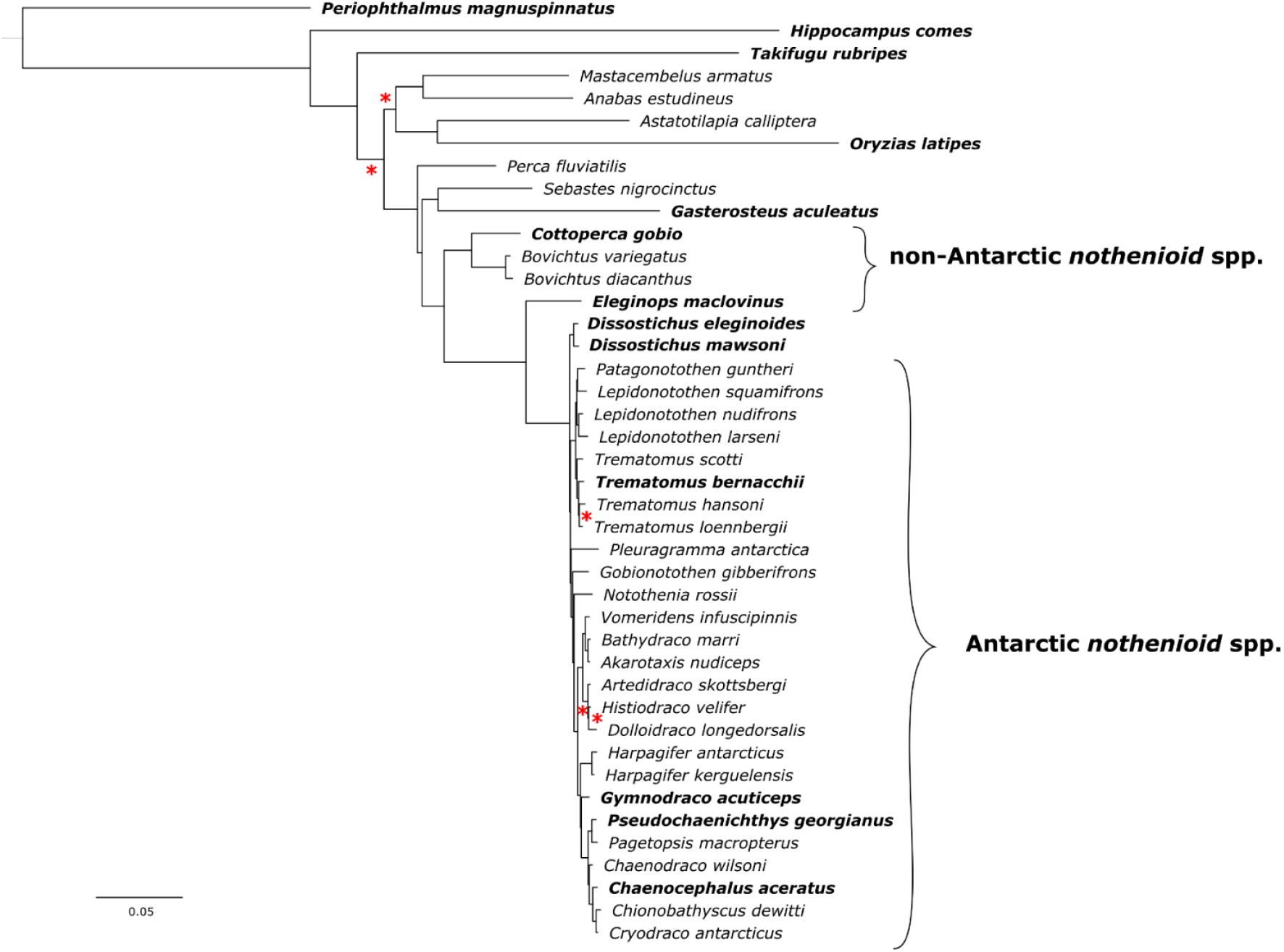
A partitioned, maximum likelihood phylogenetic analysis of 42 percomorph fish species, including 32 species of notothenioids and 10 outgroups, generated with IQ-TREE. Equal branch lengths, but different rates of evolution, were used for each one of 151 initial nucleotide partitions. A relaxed hierarchical clustering algorithm was used to examine the top ten percent of partition merging schemes and identify the best model. An ultrafast bootstrapping approach was used with 1,000 replicates. Any clades with less than 95 percent support are marked with a red asterisk. Species for which gene annotations were available are highlighted in bold.

### Assembly of the mitochondria

Assembly of the mitochondrial genome using a combination of PacBio and Illumina sequences resulted in a circular genome of 19,459 bp in length. Several duplicated genes were found, which included various deleterious mutations. The gene order in the Patagonian toothfish mitochondrial genome was quite different in comparison to other vertebrate species, as was previously described by Papetti *et al*., 2021. However, the genome we assembled had an additional pseudogene for *nad6* which was not present in the previously published Patagonian toothfish mitochondrial genome (see Figure 4 and NC_018135.1), suggesting multiple haplotypes may exist within Patagonian toothfish populations.

**Figure 4:**
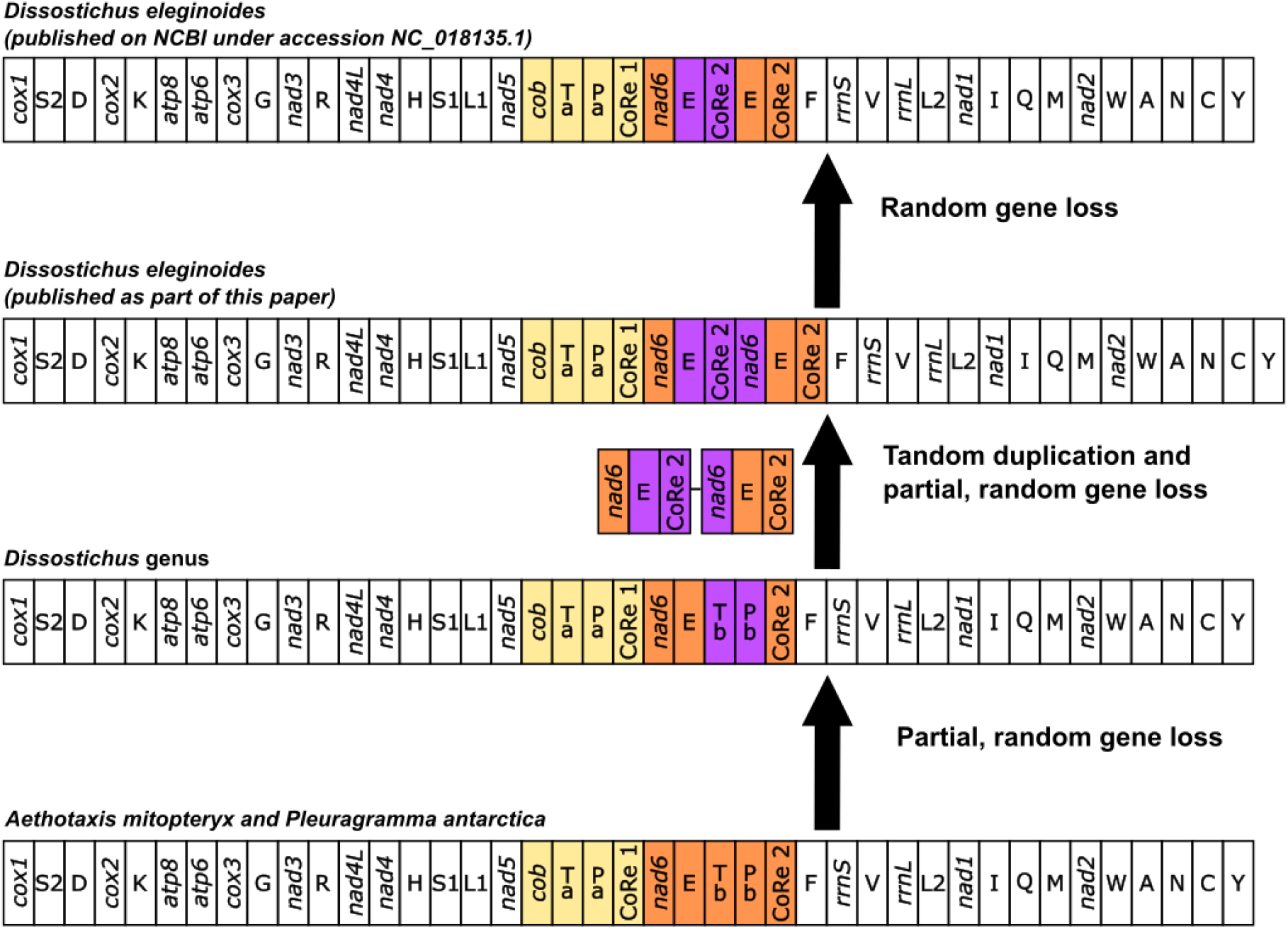
An updated proposal for mitochondrial gene order evolution based on existing work by Papetti *et al*, 2021. Gene order is linearised starting from *cox1*. Genes transposed / duplicated with respect to the gene order expected for a mitochondrial genome from a *‘standard’* vertebrate organism are shown in a yellow (gene belonging to the 5’ duplicated block) and orange (gene belonging to the 3’ duplicated block) background. Copies of the genes partially lost during the genomic rearrangement are framed in purple. Nomenclature: *atp6* and *atp8*, ATP synthase subunits 6 and 8; *cob*, apocytochrome b; *cox1-3*, cytochrome c oxidase subunits 1-3; *nad1-6* and *nad4L*, NADH dehydrogenase subunits 1-6 and 4 L; *rrnS* and *rrnL*, small and large subunit ribosomal RNA (rRNA) genes; X, transfer RNA (tRNA) genes, where X is the one-letter abbreviation of the corresponding amino acid; CoRe, Control Region.

### Comparative Genetics

Orthology analysis was carried out for every species in the phylogenetic analysis whose genome had been published with a set of gene annotations as well as *Notothenia coriiceps* (GCF_000735185.1), the closest relative of *Notothenia rossii*, for which only unannotated genomes were available. Overall, 27,297 orthogroups were identified in total, of which there were between 17,000 and 21,000 orthogroups found per species and a set of 8,442 core genes which were shared across every species. Most of the species included in the analysis had over 94 % of genes assigned to a specific orthogroup, with only *Chaenocephalus aceratus, D. eleginoides* and *Eleginops maclovinus* having a lower number of genes identified as orthologs (80-85 %). Whether differences in the latter were due to a different approach to gene annotation, or to gaps in taxonomic coverage leading to less sensitive identification of orthogroups within this species, or the species adapting to a different evolutionary niche, is unknown.

### Analysing the antifreeze glycoprotein locus

Alignment of RNA reads against each transcript predicted for haplotype 1 of the antifreeze glycoprotein (AFGP) locus (see HQ447059.1) resulted in complete coverage for trypsinogen, trypsinogen-like proteases and translocate gene coding transcripts in multiple tissues. Around 20-30% coverage was observed for transcripts coding for the chimeric antifreeze glycoprotein / trypsinogen-like protease, however such coverage was only observed for regions of the gene sharing homology with the trypsinogen-like protease gene, and not for the part of the transcript conferring the antifreeze phenotype. None of the RNA reads aligned well against any of the transcripts coding for the AFGP genes.

After using a range of different criteria to search for the AFGP locus, two candidates were identified, one from the primary assembly, and another which had previously been identified as a haplotig. The candidate loci were the only ones which aligned against more than 1000bp, with more than 90 percent identity against the AFGP haplotypes published for *D. mawsoni* (see HQ447059.1 and HQ447060.1). The candidate locus from the primary assembly was also the only one to include orthologs for the hsl and tomm40 genes which were found across different species in the notothenioid spp. clade, and where the corresponding genome regions included other features associated with the AFGP locus, such as protease and trypsin genes, as well as AFGP genes, in the *P. georgianus* genome.

To check for misassemblies PacBio sequencing reads were aligned against the candidate loci for *D. eleginoides*. The results suggested that each of the candidate loci had between 33 and 39x coverage, with a mean mapping quality of greater than 55. Manual inspection of the alignment results showed many reads which mapped across regions of the genome where additional AFGP genes were observed in other species, with no obvious signs of misassembly, insertions or deletions within the candidate loci.

Figure 5 shows a schematic represtentation of the published AFGP loci from *D. eleginoides, D. mawsoni* and *C. gobio*, indicating regions of high similarity across the toothfish loci. The *tryp3, tryp1* and *tlp* genes appear to have been duplicated in the *Dissostichus* genus relative to *C. gobio*. The *ddx6, tmen145* and *cbl* are not consistently observed across every species, though there could possibly be variation in the level of completeness of gene annotation in each of the three species. The most notable difference between the two species of *Dissostichus* is the complete absence of any AFGP gene within the AFGP locus for *D. eleginoides*. There is, however, a region found in each copy of the AFGP gene from *D. mawsoni* which shares some homology with a *tlp* gene from *D. eleginoides*. Comparing the two species of *Dissostichus* it is considered that the AFGP genes would have to have been gained, or lost, somewhere in the intergenic space between *tlp* and *cbl* genes, with both of these gene annotations being suggested to be functional, based on RNA sequencing data.

**Figure 5:**
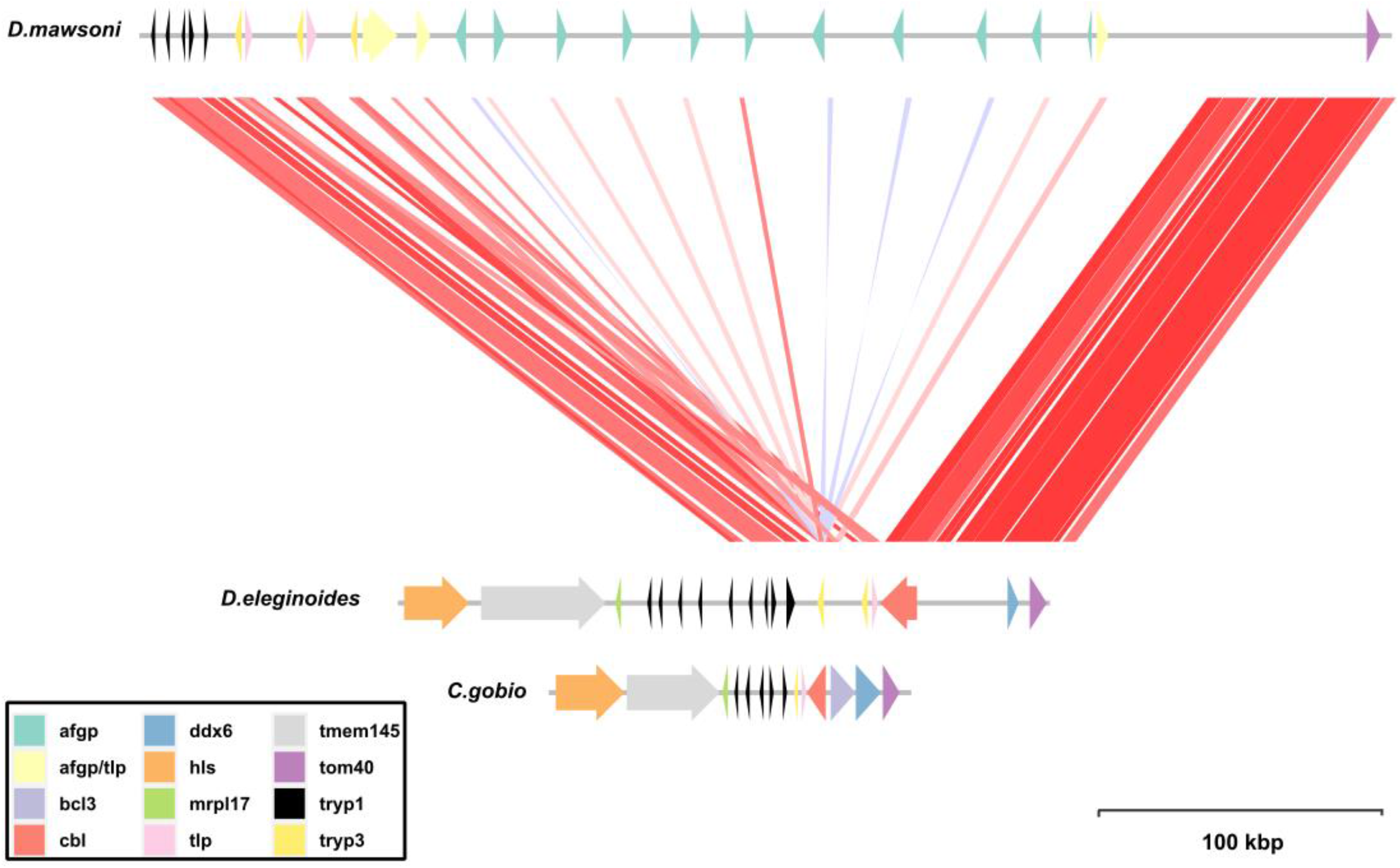
A map of the antifreeze glycoprotein (AFGP) locus for *Dissostichus eleginoides, Cottoperca gobio*, and *Dissostichus mawsoni*. Nucleotide BLAST alignments between Patagonian and Antarctic toothfish with more than 90 percent identity and a score of greater than 2830 are shown in colours ranging from red (90%) to blue (100%), which represent alignments with a high and low percentage identity. Arrow colours correspond to conserved genes; arrow heads indicate gene orientation.

## Discussion

There are many criteria for evaluating the quality of a genome, with the Vertebrate Genomes Project (VGP) having recently defined several metrics designed to assess continuity, base pair accuracy, functional completeness and chromosome status (Rhie *et al*., 2021). The genome for *D. eleginoides* presented here has been sequenced, assembled and published whilst considering a range of different criteria. The outcome is a base pair quality of greater than 40, a k-mer and BUSCO completeness score which were both higher than 90%, and an N50 value of more than 1Mb. The extensive RNA sequencing data from multiple tissues using both short and long reads, can be considered a useful resource in its own right, and be used to more effectively annotate genes, identify splice variants and confirm predictions made using *de novo* prediction algorithms (Edwards and Holt, 2013; Mudge and Harrow, 2016; Donath *et al*., 2019). The genome measures well when compared against a range of criteria set by the VGP, though one limitation of our sequencing project is that whilst the Omni-C protocol was useful for improving the contiguity of the assembly, it was not sufficient to achieve chromosomal level scaffolding nor haplotype phasing, both of which would have been made easier by combining the latest assembly algorithms with a newer generation of long read sequencing technology such as PacBio’s HiFi reads (Nurk *et al*., 2020). Depending on gene density, assemblies with a minimum N50 value of between 0.2 and 1 Mb have been shown to be sufficient to yield consistent results when being used for synteny analysis (Liu, Hunt and Tsai, 2018). This suggests that contiguity in this assembly is at least high enough to support investigation of the evolution of long, repetitive loci, like the AFGP locus (Nicodemus-Johnson *et al*., 2011), or understanding the consequences of there being different karyotypes in the two otherwise closely related species of toothfish (Ghigliotti *et al*., 2007).

In this study, we included a few comparisons across species intended to build upon earlier work, including a phylogenetic analysis based on a large number of orthologous, single copy genes from the nuclear genome, an updated look at unusual variations in the order of mitochondrial genes observed within the *Dissostichus* genus (Papetti *et al*., 2021), and comparing the order and number of genes found within the AFGP locus (Bista *et al*., 2022). The comparisons benefit from highly contiguous assemblies with good base pair accuracy and gene annotations. There is however some variation in the quality of genomes observed across the different notothenioid species (Figure 3), where only 6 out of 32 genomes submitted to various sequencing repositories appeared to include any published gene annotations, making phylogenetic analysis, identification of orthologs, and other comparisons across species challenging. In addition to missing annotations, previous studies have also reported variation in the level of completeness calculated for the genome assemblies of different notothenioid species, with gene annotations provided for the *Notothenia corriiceps, C. aceratus, E. maclovinus, D. mawsoni* assemblies having BUSCO completeness scores ranging between 80 and 97% (Bargelloni *et al*., 2019). A related issue is the level of contiguity of assemblies, with one relevant example being the first genome published for Antarctic toothfish (Chen *et al*., 2019), a species whose estimated genome size is similar to that of the Patagonian toothfish, but for which the generated genome scaffolds are much smaller in length (e.g. N90 values of 202.7 kb vs 900.4 kb for Antarctic and Patagonian toothfish, respectively). As improvements in sequencing chemistry, library preparation and assembly methods are released and standardised, it will allow the production of higher quality assemblies, and streamlined comparisons across species.

The majority of toothfish stocks are managed under the Convention for the Conservation of Antarctic Marine Living Resources (CCAMLR). It is crucial to understand the spatial stock structure of the species to enable precautionary, sustainable management. Multiple studies have been carried out to identify stock structure of both toothfish species, including analysis of tagged fish movements (Soeffker et. al. 2022, Grilly et al, 2021), mineral deposits within otoliths (Ashford *et al*., 2008), microsatellites (Smith and McVeagh, 2000) and SNP/RAD-Seq phylogenetic markers (Arkhipkin *et al*., 2022; Maschette *et al*., 2019). The Patagonian toothfish genome sequences provided by this study will further strengthen the resources available for this area of applied research, allowing identification of the most suitable restriction enzymes to use for RAD-Seq analysis (Arnold *et al*., 2013; Lepais and Weir, 2014), guiding the choice of neutral markers or restriction enzymes based on proximity to gene coding regions (Luikart *et al*., 2003) and allowing a scan of the genome to identify regions showing higher levels of adaptive or balancing selection (Beaumont and Balding, 2004; Cornman *et al*., 2013).

Within the existing literature, phylogenetic analysis of species in the notothenioids clade has been carried out using a range of techniques, including RAD-Seq (Near *et al*., 2018; Ceballos *et al*., 2019), small sets of nuclear or mitochondrial markers (Bargelloni *et al*., 1994; Matschiner, Hanel and Salzburger, 2011; Rutschmann *et al*., 2011; Near *et al*., 2012; Colombo *et al*., 2015; Maschette *et al*., 2019), as well as those based on a more comprehensive set of nuclear markers (Bista *et al*., 2022). Balushkin *et al*., (2000) proposed a single clade based on morphological criteria, which included *Pleurogramma antarctica*, the two species of *Dissostichus, Aethotaxis mitopteryx* and *Gvozdarus svetovidovi*. However, recent studies using genetic data have often identified this group as being paraphyletic (Bista *et al*., 2022), with *Pleurogramma antarctica* as an outgroup, but overall there has been insufficient evidence to reject the monophyletic hypothesis (Near *et al*., 2018) or the proposed lineages were supported by weak bootstrap values (Rutschmann *et al*., 2011; Colombo *et al*., 2015). Adding to the uncertainty, none of the analyses based on molecular evidence appear to include all of the species within the proposed group, with *Aethotaxis mitopteryx* and *Gvozdarus svetovidovi*, among others, often not being included in the analyses. Comprehensive phylogenetic analyses using genome-wide sequence information, e.g. using a large number of gene orthologs (this study), nuclear markers (Bista *et al*., 2022) or SNPs (Near *et al*., 2018), suggest that these species are very close to the base of the Antarctic clade of notothenioids, though which species is closest to the base (*Pleurogramma antarctica* (Near *et al*., 2018), *Dissostichus* spp. (Bista *et al*., 2022) or another species from the same group) still seems to be subject to some degree of disagreement, and merits further investigation.

Analysis of the transcriptome and genome of Patagonian toothfish confirmed earlier work which found no evidence for the presence of AFGP genes within the genome (Christina Cheng and William Detrich, 2007), nor expression of proteins within the blood (Gon and Heemstra, 1990). Notothenioid species lacking the antifreeze glycoprotein phenotype appear to either express AFGP, but at very low levels and with mutations in key amino acid motifs (e.g. *Notothenia angustata* and *Notothenia microlepidota*; Cheng *et al*., 2003) or there they do not express AFGP and lack the AFGP locus in their genomes (e.g. *Patagonotothen tessellata, Patagonotothen ramsayi*, and *D. eleginoides*; Cheng *et al*., 2003; Christina Cheng and William Detrich, 2007). The absence of AFGP within Patagonian toothfish suggests that either the species diverged prior to acquisition of the AFGP genotype within the nototheniid clade, the gene became degraded and was subjected to large-scale mutations, or it was lost after the fish occupied ecological niches outside of the colder waters of the Antarctic. The data generated in our study allowed for a much more detailed analysis of gene content within the AFGP locus than was possible with earlier work based on Southern blot analysis and provided no evidence for any degraded or mutated form of AFGP genes within the Patagonian toothfish genome. Unlike degeneration or mutation of the AFGP genes, it is not possible to rule out the possibility that AFGP genes were present within a common ancestor of Patagonian and Antarctic toothfish and subsequently lost. In contemporary notothenioids, a high number of copies of the gene seems to be required to survive colder habits (Cheng and Detrich, 2007; Bista *et al*., 2022), as is observed for Antarctic toothfish, for example. It is plausible that lower levels of expression of AFGP, and therefore fewer copies of the AFGP gene, would have been required for fish exploiting a slightly warmer, but still cold Southern Ocean of the recent geological past. The geological evidence suggests a gradual cooling of the climate over millions of years, with glaciers first forming in Antarctica around 35 Ma (Lagabrielle *et al*., 2009; Villa *et al*., 2013), temperatures in the Southern Ocean falling another 6 – 7°C around 14 Ma (Shevenell, Kennett and Lea, 2004) with signs of more recent cooling within the last few million years (Bista *et al*., 2022). As already discussed, more recent phylogenetic analyses suggest the Patagonian toothfish diverged from the rest of the notothenioids clade relatively early on, when temperatures in the Antarctic were probably warmer than they are at present, leaving open the possibility that its AFGP locus could indicate an earlier state, prior to large scale duplication of the AFGP gene which subsequently led to the high expression of AFGP seen in other species. Reconstructing the evolution of genes within this locus with any degree of confidence is likely to require analysis of more examples from members of the Pleuragrammatinae clade, as well as other species thought to be lacking copies of the AFGP gene.

## Conclusion

In this study, we produced a high-quality genome assembly for the Patagonian toothfish, an ecologically and economically important fish in the sub-Antarctic regions of the Southern Ocean. A second haplotype for the Patagonian toothfish mitochondrial genome was identified, with the potential for both haplotypes to be used as population markers if they can be shown to exist within specific populations. Predicted gene sequences, together with the transcriptomic data generated for a variety of tissues in this study, will facilitate studies on physiology, disease, reproduction and population genetics in this species. Our work found no evidence of the presence of AFGP genes in the Patagonian toothfish genome. Phylogenetic analysis based on a set of orthologous protein sequences showed that the Patagonian toothfish is near the root of the notothenioids clade. The genome will provide a valuable genetic resource for physiological, ecological and evolutionary studies on this species.

## Acknowledgements

We thank Zoe Fowler (Department of Agriculture, Falkland Islands), Sue Gregory (GSGSSI) and observers onboard the vessel for collecting the required tissue samples, Dr Mark Belchier (GSGSSI) and several colleagues for transporting samples to the UK. We acknowledge Dr Aaron Jeffries and the Exeter Sequencing Facility for advise on the sequencing strategy and for conducting the PacBio and Illumina sequencing included in this manuscript. This work was funded by Argos Froyanes Ltd.

## Data availability

All raw data, genome assembly and annotations have been deposited to the National Center for Biotechnology Information (NCBI) databases under BioProject PRJNA864592 and Biosamples SAMN30075165 and SAMN30114550-SAMN30114559. The Whole Genome Shotgun project has been deposited at DDBJ/ENA/GenBank under the accession JAOVFM000000000. The version described in this paper is version JAOVFM010000000.

## Supplemental Material

<u>Supplemental Table 1</u> - Fish species used for orthology and phylogenetic analyses in this study.

<u>Supplemental Table 2</u> – Identified repeat elements in the Patagonian toothfish genome

<u>Supplemental Table 3</u> - Lists of enriched gene ontologies as determined by Gene Set Enrichment Analysis (GSEA) on lists of genes expressed in each individual tissue in comparison to genes expressed all other tissues together.

